# A CXCL12 morphogen gradient uncovers lung endothelial heterogeneity and promotes distal vascular growth

**DOI:** 10.1101/2022.04.30.490096

**Authors:** Prashant Chandrasekaran, Nicholas M. Negretti, Aravind Sivakumar, Maureen Peers de Nieuburgh, Joanna Wang, Nigel S. Michki, Fatima N. Chaudhry, Hongbo Wen, Sukhmani Kaur, MinQi Lu, Jarod A. Zepp, Lisa R. Young, Jennifer M.S. Sucre, David B. Frank

**Author notes:** Corresponding Authors David B. Frank, M.D., Ph.D., Children’s Hospital of Philadelphia, 3615 Civic Center Blvd, ARC 416 K, Philadelphia, PA 19104, Jennifer M.S. Sucre, M.D. Department of Pediatrics, Vanderbilt University, 2215 Garland Ave, 1125 Light Hall, Nashville, TN 37232. Equal contribution (Prashant Chandrasekaran, Nicholas M. Negretti, Aravind Sivakumar). Equal contribution (Jennifer M.S. Sucre, David B. Frank).

## Abstract

In adults, there is a growing amount of data uncovering the cellular diversity of the pulmonary circulation and mechanisms governing vascular repair after injury, however, molecular and cellular mechanisms contributing to the morphogenesis and growth of the pulmonary vasculature during embryonic development are less clear. Importantly, deficits in vascular development lead to a large number of lung diseases in children, indicating a need to uncover fetal programs that promote pulmonary vascular growth. To address this, we used a transgenic mouse reporter for expression of *Cxcl12*, an arterial hallmark gene, and performed single-cell RNA sequencing on isolated *Cxcl12*-DsRed+ endothelium to assess cellular heterogeneity within pulmonary endothelium. Combining cell annotation, gene ontology analysis, and spatial transcriptomics allowed us to segregate the developing artery into spatially and functionally distinct novel subpopulations. In addition, expression of *Cxcl12* suggests a morphogen gradient from arteries to capillaries, suggesting directed cell migration for pulmonary vascular development. Disruption of this gradient led to abnormal branching and pulmonary vascular hypoplasia. These data provide evidence for arterial endothelial functional heterogeneity and reveal conserved signaling mechanisms essential for pulmonary vascular development.

## INTRODUCTION

Transition of the blood through the lungs for respiration is dependent on an intact and adequate pulmonary vascular tree. Disruption due to prematurity, congenital malformations and injury results in a deficient vascular bed leading to sequelae such as edema, hypoxemia, and pulmonary hypertension. Prevention of the long-term effects of these sequelae depends on efficient regeneration and re-initiation of vascular fetal programs to restore the circulation and preserve gas exchange (Cao et al., 2016; Ding et al., 2011; Mammoto et al., 2019; McCulley et al., 2018; Rafii et al., 2015; Vila Ellis et al., 2020; Zhao et al., 2020). This coordinated repair effort is critical in preventing significant morbidity and mortality. As such, studies identifying the cellular and molecular mechanisms of pulmonary vascular development may inform regeneration and/or re-initiation of development of the vasculature following injury.

Growing evidence indicates that the endothelium of the pulmonary circulation is heterogeneous. Studies using single-cell RNA sequencing analysis on the developing and adult lung have uncovered distinct endothelial subpopulations comprising the arteries, veins, capillaries, and lymphatics (Asosingh et al., 2021; Gillich et al., 2020; Guo et al., 2019; He et al., 2018; Negretti et al., 2021; Niethamer et al., 2020; Rodor et al., 2021; Saygin et al., 2020; Schupp et al., 2021; Travaglini et al., 2020; Vila Ellis et al., 2020; Zepp et al., 2021). In addition, insights from these studies suggest functional specialization of endothelium. In particular, alveolar capillaries are composed of two populations predicted to participate in respiration or regeneration after injury based on unique transcriptional profiles (Gillich et al., 2020; Niethamer et al., 2020; Vila Ellis et al., 2020). Unlike capillary endothelium, data are limited in the macrovasculature during development and adult disease and homeostasis (Asosingh et al., 2021). To date, there are no data profiling subpopulations of arterial endothelium in the developing lung. Identification and characterization of distinct functional arterial endothelial cell subtypes and their signaling mechanisms during development is necessary for understanding the processes of growth, branching, angiogenesis, and vessel assembly for tissue regeneration and replacement.

To assess arterial heterogeneity in the lung, we isolated arterial endothelium based on a previously identified arterial marker, C-X-C motif Chemokine12 (CXCL12). CXCL12 promotes growth and chemoattraction of cell expressing one of two receptors, C-X-C Chemokine Receptor 4 (CXCR4), and an alternative receptor, Atypical Chemokine Receptor 3 (ACKR3) (Kiefer and Siekmann, 2011). CXCL12 plays a critical role in tissue vascularization (Das et al., 2019; Ivins et al., 2015; Tachibana et al., 1998; Takabatake et al., 2009; Xu et al., 2014), and deletion of CXCL12 or its receptors leads to abnormal pulmonary artery branching in development and impaired capillary regeneration in a pneumonectomy model in adult mice (Kim et al., 2017; Rafii et al., 2015). Using fluorescence activated cell sorting (FACS) of endothelial cells (ECs) from lungs of a *Cxcl12*^*DsRed*^ reporter mouse (Ding and Morrison, 2013), we performed single-cell RNA sequencing on 26,652 cells across development. Surprisingly, we found a *Cxcl12* expression gradient marking not only macrovascular arterial ECs (maECs), but also the subsets of previously identified capillary ECs with high and low levels of *Cxcl12* expression, respectively. Using unbiased clustering and cell annotation, we observed proximal and distal patterning of the arterial endothelium across development with unique gene enrichment consisting of genes associated with vascular wall assembly and angiogenesis, respectively. Finally, deletion of CXCL12 during development resulted in not only previously identified aberrant arterial branching but also pulmonary vascular hypoplasia, indicating a critical role for CXCL12 signaling axis in pulmonary vascular assembly and development.

## RESULTS

### Morphometric analysis of the developing pulmonary arterial system

We performed an imaging analysis of the growing arterial tree throughout development to identify periods of rapid vascular growth and significant morphometrical changes. Using the *Cxcl12*^*DsRed*^ reporter mouse with *DsRed* knocked into exon 1 of the *Cxcl12* locus, we acquired whole lung or single lobe imaging of the developing arteries at E12.5, E13.5, E15.5, and E18.5 using confocal and Leica Thunder microscopy systems (Fig. 1A-E). At several stages, we confirmed specificity of the reporter using immunohistochemistry (IHC) for DsRed reporter expression and markers for endothelium including ETS related gene (ERG) and endomucin (EMCN). As in other organ systems, *Cxcl12*-DsRed is expressed highly throughout the arterial endothelium with additional expression observed in vascular smooth muscle cells (VSMCs) (Fig S1A-F) (Chang et al., 2017; Ghadge et al., 2021; Ivins et al., 2015; Takabatake et al., 2009). We observed no DsRed expression in vein endothelium during these developmental time periods. As early as E12.5, DsRed was expressed at high levels in the pulmonary arteries as two primary intrapulmonary branches (Fig. 1A). One day later, we observed the initiation of secondary branching from the primary intrapulmonary artery with small, thin, tubular structures appearing to sprout from the intrapulmonary artery and grow outward to the lung periphery (Fig. 1B,C). By E15.5, the intrapulmonary artery evolved with the formation of additional secondary branches along with tertiary and quaternary arterial branches (Fig. 1D). Following this, the arterial tree underwent significant growth and further branching complexity by the end of prenatal vascular development (Fig. 1E).

**Figure 1.**
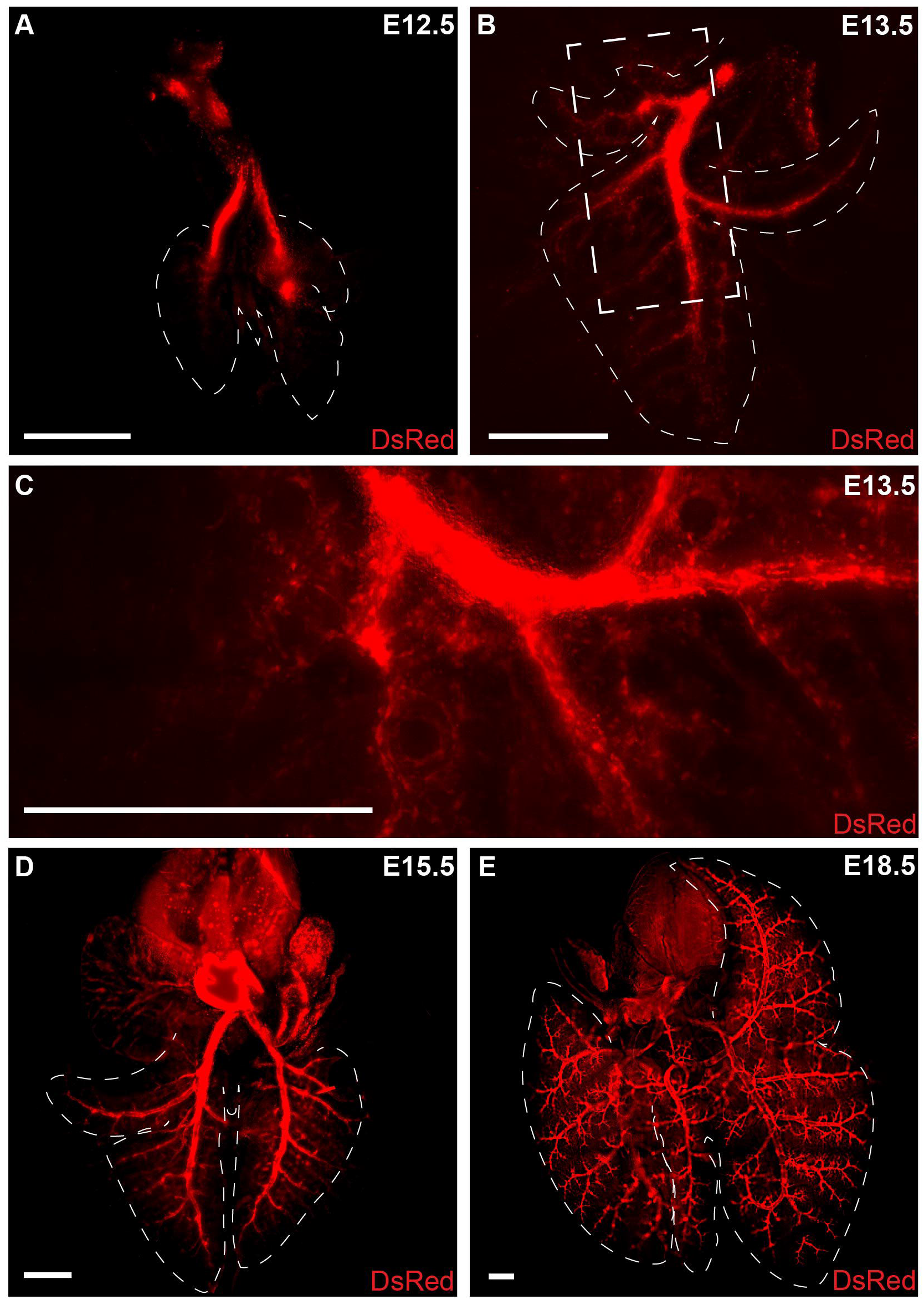
Morphogenesis of the developing pulmonary arterial tree in *Cxcl12*^*DsRed/+*^ reporter embryos. (A) E12.5, two intrapulmonary arteries are visible. (B) E13.5, Secondary branches extend out from the primary branches. (C) Higher magnification of secondary branching. (D) E15.5, increased secondary branching along with tertiary and quaternary branches. (E) E18.5, robust and complex branching is apparent. Scale bar: 500 µm.

### Single-cell transcriptomic analysis uncovers heterogeneity of the *Cxcl12*+ endothelium

Using the data from our morphometric analyses, we selected specific developmental periods with significant changes in structure and growth to assess cellular and transcriptomic composition of the growing arterial tree (Fig. 2). Single-cell suspensions were collected from 2-4 embryonic or pup combined lungs acquired at E13.5, E15.5, E18.5, and P8). We obtained *Cxcl12*-DsRed+ ECs using a FACS-based strategy (DsRed+/CD31+/EpCAM-/CD45-). Single-cell RNA sequencing (scRNA-seq) was performed using a droplet-based platform (10x Genomics Chromium). Across the four timepoints, a total of 26,652 endothelial single cell transcriptomes were analyzed, with 391 cells at E13.5, 9,082 cells at E15.5, 9016 cells at E18.5, and 8,163 cells at P8 (Fig. 2A). The median unique molecular identifiers (UMIs) per cell was 6,666 (interquartile range 6253-7545), and a median of 2,483 genes (interquartile range 2059-3003) was detected per cell.

**Figure 2.**
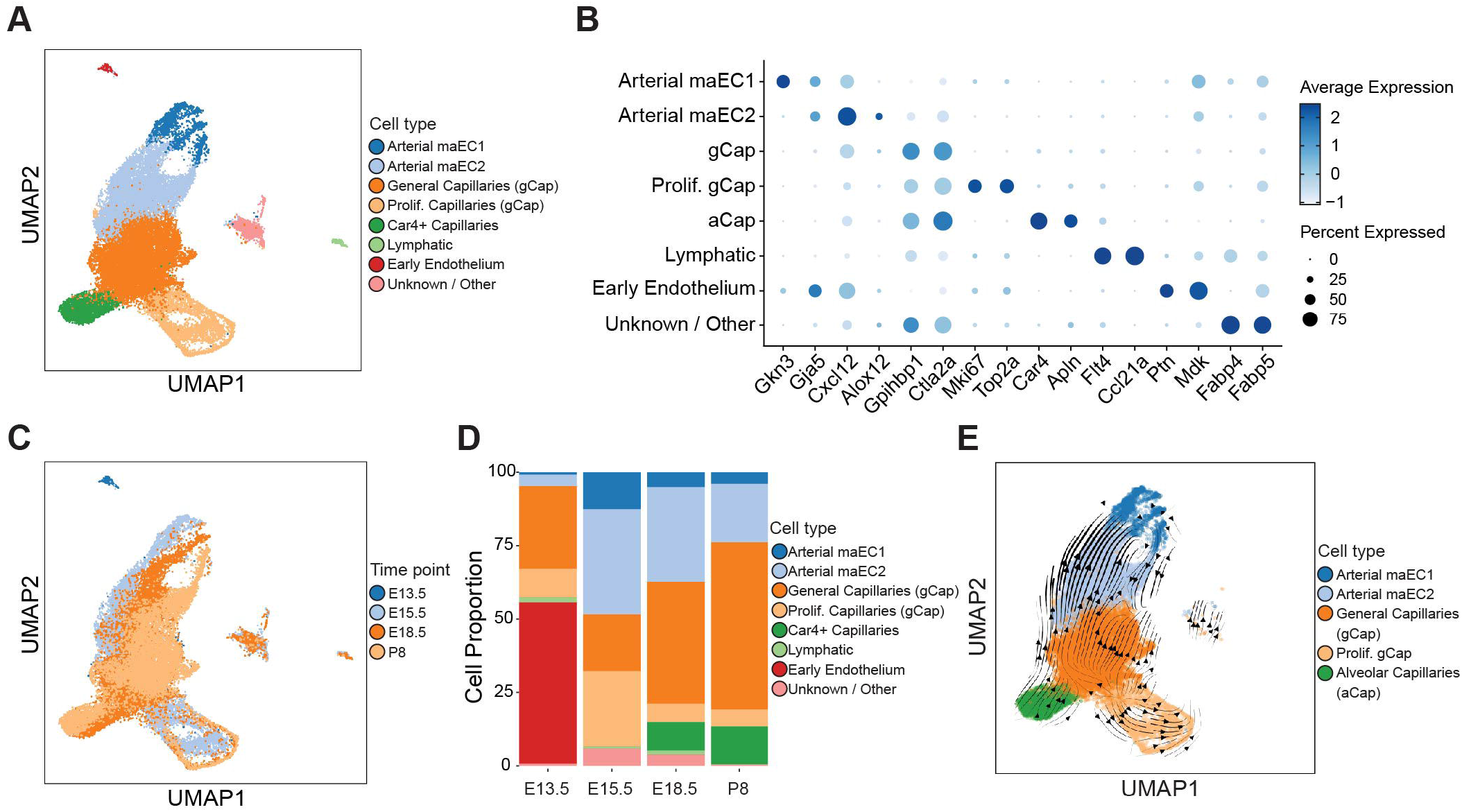
Sequencing of *Cxcl12*+ cells uncovers heterogeneity and novel arterial EC subpopulations. (A) UMAP embedding of *Cxcl12*-DsRed+ cells (n = 26,652) colored by cell type. (B) Dot plot of marker genes for each cell type where dot size indicates proportion of cells within a cluster expressing a gene, and color intensity indicates relative expression level. (C) UMAP embedding colored by time point. (D) Cellular composition changes in *Cxcl12*+ ECs over time. (E) RNA velocity vectors were calculated with CellRank and overlayed on the UMAP embedding. Thicker lines indicate greater predicted RNA velocity.

Louvain clustering was applied to the single-cell transcriptomes to identify transcriptionally related clusters, identifying 8 cellular populations including two novel arterial EC populations: Arterial maEC1 marked by *Gkn3* and *Gja5* expression, arterial maEC2 marked by elevated *Cxcl12* and *Alox12 expression*, general capillaries (gCap) marked by *Gpihbp1* and *Ctla2a* expression, proliferating gCaps marked by *Mki67* and *Top2a* expression, alveolar capillaries (aCap) marked by *Car4* and *Apln* expression, lymphatic endothelium marked by *Flt4* and *Ccl21a* expression, early endothelium marked by *Ptn* and *Mdk*, and a cluster of unknown / other endothelial cells marked by *Fabp4* and *Fabp5* expression (Fig. 2B, Table S1). Examining the UMAP by sample timepoint indicated that the E13.5 is the most distinct from the others, and that the other three timepoints (E15.5, E18.5, and P3) have relatively similar cellular composition except for the proliferating gCap cells which decrease in abundance from E15.5 to E18.5 (Fig. 2C-D). Further analysis of RNA-velocity based cellular transitions suggested potential transitions between arterial maEC1, arterial maEC2, and gCap populations (Fig. 2E). To assess the genes that share expression patterns with Cxcl12, a spearman correlation coefficient was calculated between every gene and Cxcl12. There were 23 genes with a correlation coefficient greater than 0.3, including *Gja4* and *Slc6a6* (Fig. S2).

### Temporal allocation of CXCL12 endothelium reveals distinct spatial and functional populations

Analysis of transcriptomic data identified two novel subpopulations of maECs that persist throughout development (Fig. 2A), and we have identified *Gkn3* and *Alox12* specifically defining these clusters in arterial maEC1 and arterial maEC2 subpopulations, respectively (Fig. 3A,B). GKN3 has no previously described role in lung development, but it has been identified as a possible receptor for the Japanese Encephalitis Virus in the brain (Mukherjee et al., 2018), and recently described as an arterial endothelial marker in the brain (Vanlandewijck et al., 2018). ALOX12 is an enzyme involved in arachidonic acid metabolism that promotes tumor progression and angiogenesis (Zheng et al., 2020). Importantly, ALOX12 has also been implicated in hypoxia-induced endothelial angiogenesis and smooth muscle cell proliferation in the lung (Preston et al., 2006; Zhang et al., 2018). To characterize functional differences between these two populations, we generated heatmaps to interrogate differential gene expression of the top 40 positively associated marker genes for each cluster and used Panther Gene Ontology (GO) enrichment analysis to predict potential biological functional differences based on differential gene expression for each population (Fig. 3C-E and Table S2). While there was some overlap in the functions identified, we noted several distinct roles for each group. For arterial maEC1, the top representative genes predicted roles in extracellular matrix and structure organization, external encapsulating structure organization, and multicellular organism development (Fig. 3D). *Eln, Mgp, Fbln5, Fbln2, Fn1*, and *Lox* comprised a portion of these genes and suggest a function in vessel wall assembly. Arterial maEC2 expressed many genes associated with vascular development as noted by GO analysis (Fig. 3E). Genes include *Jag1, Fbn1, Sox17, Gata6, Cxcl12, Hey1, Tgfb2*, and *Vegfc*.

**Figure 3.**
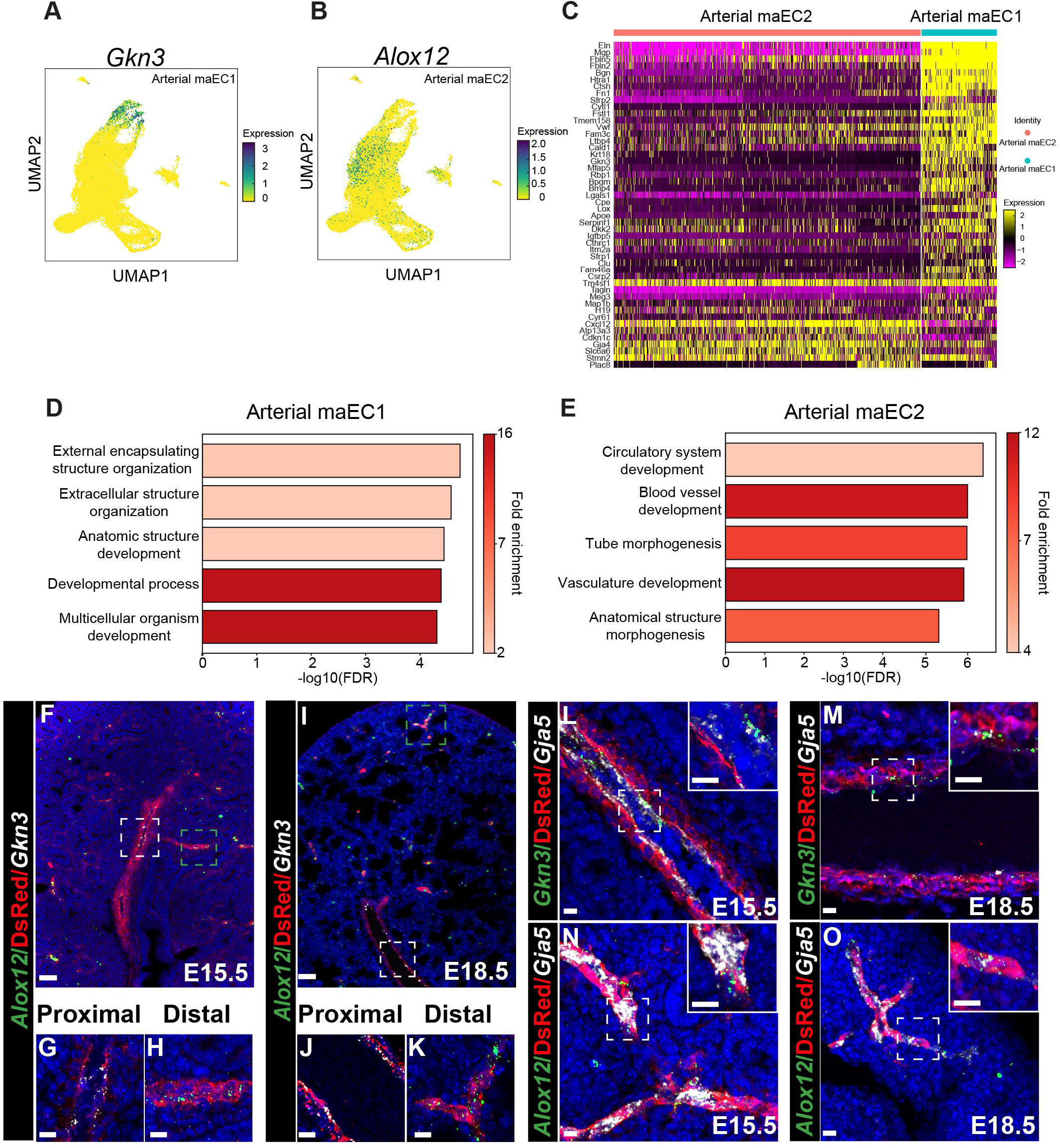
Identification of proximal and distal arterial ECs by spatial transcriptomic and predicted function analysis. (A) UMAP embedding of cell colored by *Gkn3* expression (B) UMAP embedding of cells colored by *Alox12* expression. (C) Heat map representing expression of hallmark genes in arterial macro endothelial cell types 1 and 2 (arterial maEC1 and maEC2). (D) Gene ontology (GO) analysis of maEC1 cluster (E) GO analysis of maEC2 cluster. (F-K) RNA ISH for *Gkn3* and *Alox12* and IHC for DsRed protein at E15.5 (F-H) and E18.5 (I-K) with magnification of proximal (G,J) and distal (H,K) areas. (L, M) RNA ISH with IHC of proximal lung tissue regions for *Gkn3, Gja5*, and DsRed at E15.5 (L) and E18.5 (M). (N, O) RNA ISH with IHC of distal lung tissue regions for *Alox12, Gja5*, and DsRed at E15.5 (N) and E18.5 (O). Insets: Higher magnification of boxed areas. Scale Bar: F,I = 50 µm, G,H,J,K =10 µm, L-O insets = 10 µm.

The distinct functional attributes suggested by gene expression and ontology analysis point to spatially-defined arterial endothelial compartmentalization. Given that arterial maEC2s were enriched in genes involved in vascular development and in a spatial continuum with capillary ECs on UMAP-based clustering, we hypothesized that they comprise a distal arterial maEC (distal maEC2) population. In addition, arterial maEC1 had gene enrichment in extracellular structure suggesting a functional role in vessel wall assembly. Combined with their spatial location further away from capillary ECs on UMAP-based clustering, we hypothesized that they represented a proximal arterial ma EC (proximal maEC1) population. To test this, we performed spatial transcriptomics for *Alox12* and *Gkn3* using RNA in situ hybridization (RNA ISH) combined with IHC for DsRed in *Cxcl12*^*DsRed*^ reporter embryonic lung tissue at E15.5 and E18.5. At both time points, *Alox12* localized to the distal arterial endothelium with no expression in the more proximal arteries (Fig. 3F-K). By contrast, *Gkn3* is expressed proximally with extension limited peripherally. Both distal Alox12+ and proximal Gkn3+ arterial EC populations colocalized with *Gja5*, confirming specificity to the arterial ECs (Fig. 3L-O). In addition, *Gkn3* endothelial expression accompanied expression of both smooth muscle actin (SMA) and tropoelastin (ELN) supporting a role in endothelial-mediated vessel wall assembly (Fig. S3A,B).

Interestingly, scRNA-seq of the *Cxcl12*-DsRed+ endothelium included not only arterial endothelium but also capillary endothelium (Fig. 2A). This comprised all previously described capillary EC populations including general capillary (gCap), proliferating gCap, and alveolar capillary (aCap) cells, marked by expression of *Gpihbp1, Mki67*, and *Apln* or *Car4*, respectively. Subpopulations aCap and gCap have been recently described as capillary 1 cells (CAP1), proliferating CAP1 cells, and capillary 2 cells (CAP2) (Sun et al., 2022). In concordance with previous data, in our study both gCap and aCap populations arose after E15.5 and are apparent by E18.5 (Gillich et al., 2020; Negretti et al., 2021; Vila Ellis et al., 2020; Zepp et al., 2021). We validated the transcriptional profile and localized these populations at the tissue-level with RNA ISH for *Cxcl12, Apln*, and *Gpihbp1* transcripts in combination with DsRed expression by IHC (Fig. S3C,D). Again, expression of *Cxcl12* transcript resided predominantly in the arterial endothelium. With higher magnification, we observed rare transcript expression in capillary endothelium.

### Characterization of a CXCL12 morphogen gradient and signaling axis

Low *Cxcl12* expression in the distal capillary plexus and high expression in the arterial tree indicated a CXCL12 morphogen gradient. Morphogen gradients recruit distal cells in a process of directed cell migration called haptotaxis (Majumdar et al., 2014). Haptotaxis involving CXCL12 signaling has been well studied in mammalian organ systems and organisms such as zebrafish (Dona et al., 2013; Ivins et al., 2015; Xu et al., 2014). We performed UMAP embedding of *Cxcl12*-DsRed+ ECs confirming the presence differential expression of *Cxcl12* across endothelial subpopulations (Fig. 4A and 4D). From this, we predicted a spatial CXCL12 gradient in the tissue, and we determined the EC subpopulations expressing the CXCL12 receptors, CXCR4 and ACKR3. UMAP embedding of *Cxcl12*-DsRed+ endothelium colored by *Cxcr4* expression indicated a cellular distribution to the distal maEC2s and a small portion of the capillary ECs (Fig. 4B and 4E). Similar embedding by *Ackr3* confined expression to capillary ECs (Fig. 4C and 4F). Since *Cxcl12*-DsRed + endothelium was void of venous identity in the lung, we reanalyzed previously published single-cell sequencing data on endothelium from the developing mouse lung for the CXCL12 signaling axis distribution (Negretti et al., 2021). While *Cxcr4* remained localized in arterial and capillary endothelium, *Ackr3* is expressed not only in the capillary endothelium but also robustly in venous endothelium (Fig. S4A-D) throughout prenatal and postnatal lung development. In addition, C*xcl12* becomes more broadly expressed in the developing postnatal lung and can be observed in venous endothelium (Fig. S4E,F).

**Figure 4.**
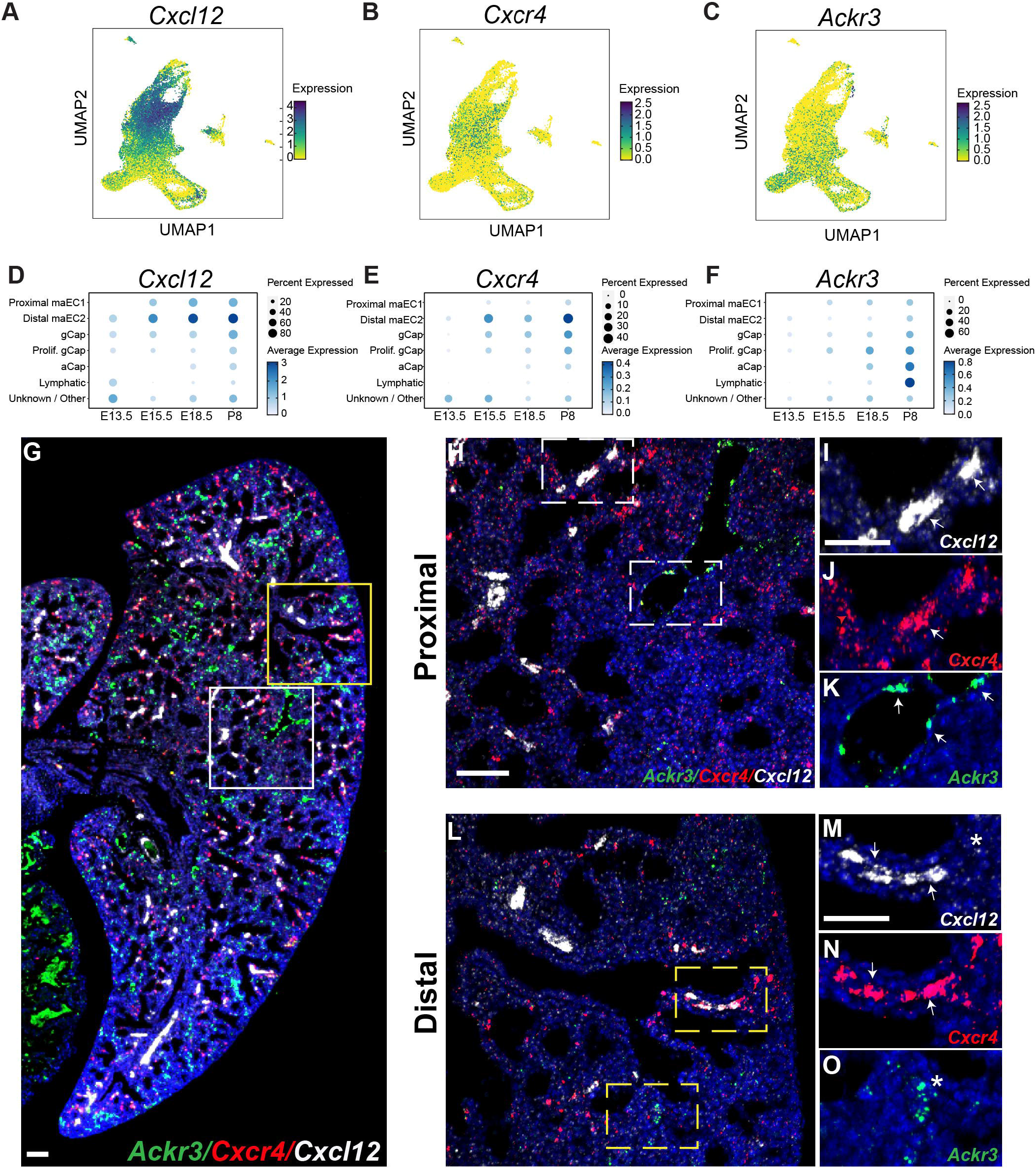
Characterization of *Cxcl12* morphogen gradient and signaling axis. (A) UMAP embedding of cells colored by *Cxcl12* expression. (B) UMAP embedding of cells colored by *Cxcr4* expression. (C) UMAP embedding of cell colored by *Ackr3* expression. (D) Dot plots of *Cxcl12* expression in different EC compartments at E13.5, E15.5, E18.5 and P8. (E) Dot plots of *Cxcr4* expression at same timepoints. (F) Dot plots of *Ackr3* expression at same timepoints. (G) RNA ISH for spatial expression of *Ackr3, Cxcr4*, and *Cxcl12* at E18.5. (H) Higher magnification of RNA ISH for *Ackr3, Cxcr4*, and *Cxcl12* in proximal lung at E18.5. (I-K) Higher magnification of boxed areas. (L) Higher magnification of RNA ISH for *Ackr3, Cxcr4*, and *Cxcl12* in proximal lung at E18.5. (M-O) Higher magnification of boxed areas. Scale: G = 100 µm, H,L = 50 µm, I-K, M-O = 50 µm. Arrows represent expression in macrovessels * represent expression in capillaries.

We performed tissue-level validation of expression of the CXCL12 signaling axis using RNA ISH on E15.5 and E18.5 C*xcl12*^*DsRed/+*^ embryonic tissue. *Cxcr4* and *Ackr3* expression was concomitantly determined with *Cxcl12* expression. *Cxcr4* is expressed in the distal maEC2s overlapping with robust *Cxcl12* expression (Fig. 4G-O). In addition, a scattering of capillary ECs contained *Cxcr4* (Fig. 4J and 4N). Similarly, *Ackr3* localized to capillary ECs, but it was more prominent in the venous endothelium (Fig. 4K and 4O). Ackr3 and Cxcr4 were not colocalized in the distal vasculature. Again, *Cxcl12* transcript and DsRed reporter protein expression were highest and prevalent mostly in arterial endothelium only. While there was significant overlap of *Cxcr4* and *Cxcl12* in the distal maEC2s, *Cxcl12* and *Ackr3* shared far less co-localization with *Ackr3* almost exclusively in the capillary and vein endothelium (Fig. S4H-K).

### Global loss of CXCL12 results in distal branching defects and pulmonary vascular hypoplasia

Expression and localization of the CXCL12 signaling axis indicated a morphogen gradient for directional migration and formation of the distal vasculature. To disrupt this gradient, we generated *Cxcl12* heterozygous (*Cxcl12*^*DsRed/+*^) controls and *Cxcl12* homozygous null (*Cxcl12*^*DsRed/DsRed*^) embryos. As previously reported, no postnatal *Cxcl12*^*DsRed/DsRed*^ were recovered (Kim et al., 2017). Whole mount immunofluorescence imaging was performed first to assess morphological changes in the microvasculature at E18.5. In concordance with previously published data, we observed proximal and distal branching defects (Kim et al., 2017) (Fig. 5A,B). In addition, we noted diminished mean vessel diameter for primary to quaternary branches in null embryos (Fig. 5C) Distally, the arteries lacked directionality toward the periphery that is preserved in control embryos (Fig. S5A,B). Quantification of branching angles of the distal arteries using Filament Tracing in Imaris revealed discrepant angles in *Cxcl12*^*DsRed/DsRed*^ embryos compared to controls (Fig. S5C). Additionally, Sholl analysis that measures branching complexity via quantification of vessel intersections of drawn concentric shells from the main branch was abnormally high compared in *Cxcl12*^*DsRed/DsRed*^ embryos compared to controls (Sholl, 1953) (Fig. S5D,E). Furthermore, *Cxcl12*^*DsRed/DsRed*^ embryonic lungs appeared to have an increase in total branching number, indicating a compensatory mechanism to increase vessel density (Fig. 5D-H).

**Figure 5.**
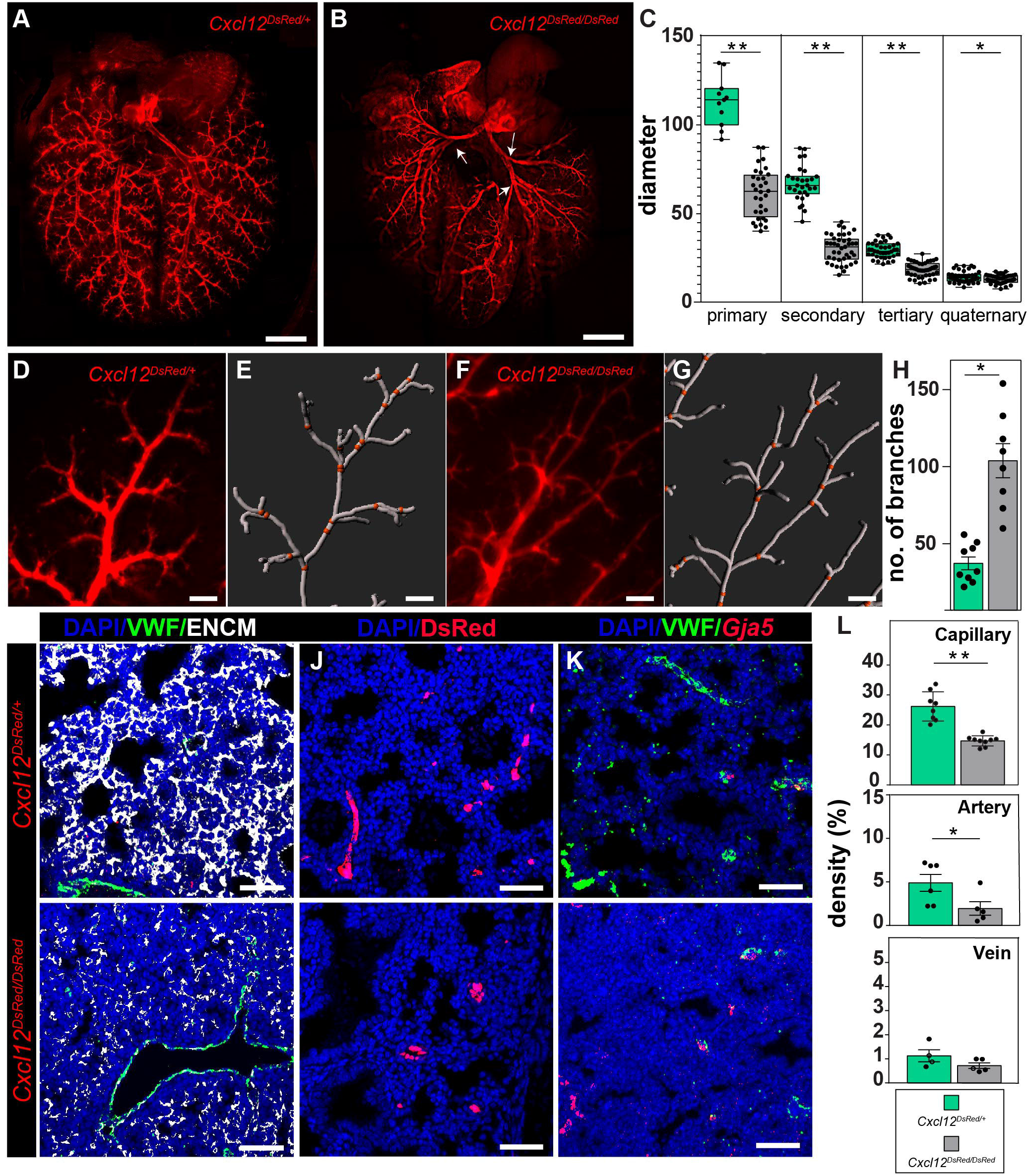
Loss of *Cxcl12* leads to branching defects and pulmonary vascular hypoplasia. (A-B) Whole mount imaging of arterial endothelium in *Cxcl12*^*DsRed/+*^ (A) and *Cxcl12*^*DsRed/DsRed*^ (B) at E18.5. *Cxcl12*^*DsRed/DsRed*^ lungs show defective proximal branching (white arrows). (C) Arterial diameter distribution of control *Cxcl12*^*DsRed*^ versus null *Cxcl12*^*DsRed/DsRed*^ arteries (n=4, filaments measured ≥ 10, Statistical significance **p<0.005, *p<0.05). (D-H) Branching analysis using Imaris Filament analysis. (D,E) Representative image of *Cxcl12*^*DsRed/+*^ artery and Filament tracing using IMARIS 9.2. (F, G) Representative image of *Cxcl12*^*DsRed/DsRed*^ artery and Filament tracing. (H) Number of branches from the conserved motif regions in the periphery (n=4, images analyzed ≥ 8, Statistical significance: *p<0.05). (I-L). Vascular density measurements in control *Cxcl12*^*DsRed*^ (top panels) versus null *Cxcl12*^*DsRed/DsRed*^ (lower panels) lung tissue. (I) Capillary density IHC staining of tissue sections for VWF (macrovessel) and ENCM (capillary and vein). (J) Arterial density IHC staining of tissue sections for DsRed (artery). (K) Venous density IHC and RNA ISH staining for *Gja5* (artery) VWF (macrovessel) expression. (L) Volume density calculations of capillaries, arteries and veins (n≥4, **p<0.005, *p<0.05. Scale: A-B = 1mm, D-G = 100 µm, I-K = 50 µm.

CXCL12 and its receptors, CXCR4 and ACKR3, were expressed in a compartmental fashion. Thus, deficits in the vascular bed may expand beyond the arterial endothelium. To quantify arterial, vein, and capillary endothelium, we performed IHC and RNA ISH for multiple combinations of markers of arterial, vein, and capillary endothelium. We used von Willebrand factor (VWF) to identify macrovessels, *Gja5* and DsRed for arterial endothelium, neuregulin-1 (NRG1) for vein endothelium, and EMCN for capillary and vein endothelium on lung tissue sections at E18.5 (Fig. 5I-L and S5F-I). Using three-dimensional (3D) surface rendering, we were able to quantify the vessel density in tissue sections by calculating percent of volume of capillary, artery, or vein in each section (Chen et al., 2021; Nizari et al., 2019). Capillaries comprised a majority of the vessel density, and capillary density was significantly reduced in the *Cxcl12*^*DsRed/DsRed*^ embryos compared to controls (Fig. 5I,L and S5F). Similarly, arterial density was modestly reduced, but the density of veins was unchanged (Fig. 5J-L and S5G,H). We also calculated the total number of macrovessels in a tissue section as branching was increased in *Cxcl12*^*DsRed/DsRed*^ embryos (Fig. S5I). While there was no significant change in arterial number, we did note a significant decrease in the number of veins.

To corroborate our quantitative findings and ascertain affected cell populations, we performed scRNA-seq on whole E18.5 lungs from control and *Cxcl12*^*DsRed/DsRed*^ embryos (Fig. S6A,B). Overall, we saw that the percentage of ECs in the lung was diminished (Fig. S6C). We extracted the EC data and re-clustered to examine changes in the cellular representation of the endothelium (Fig. 6A,B). From these data, we were able to reidentify all previously referenced EC clusters.

**Figure 6.**
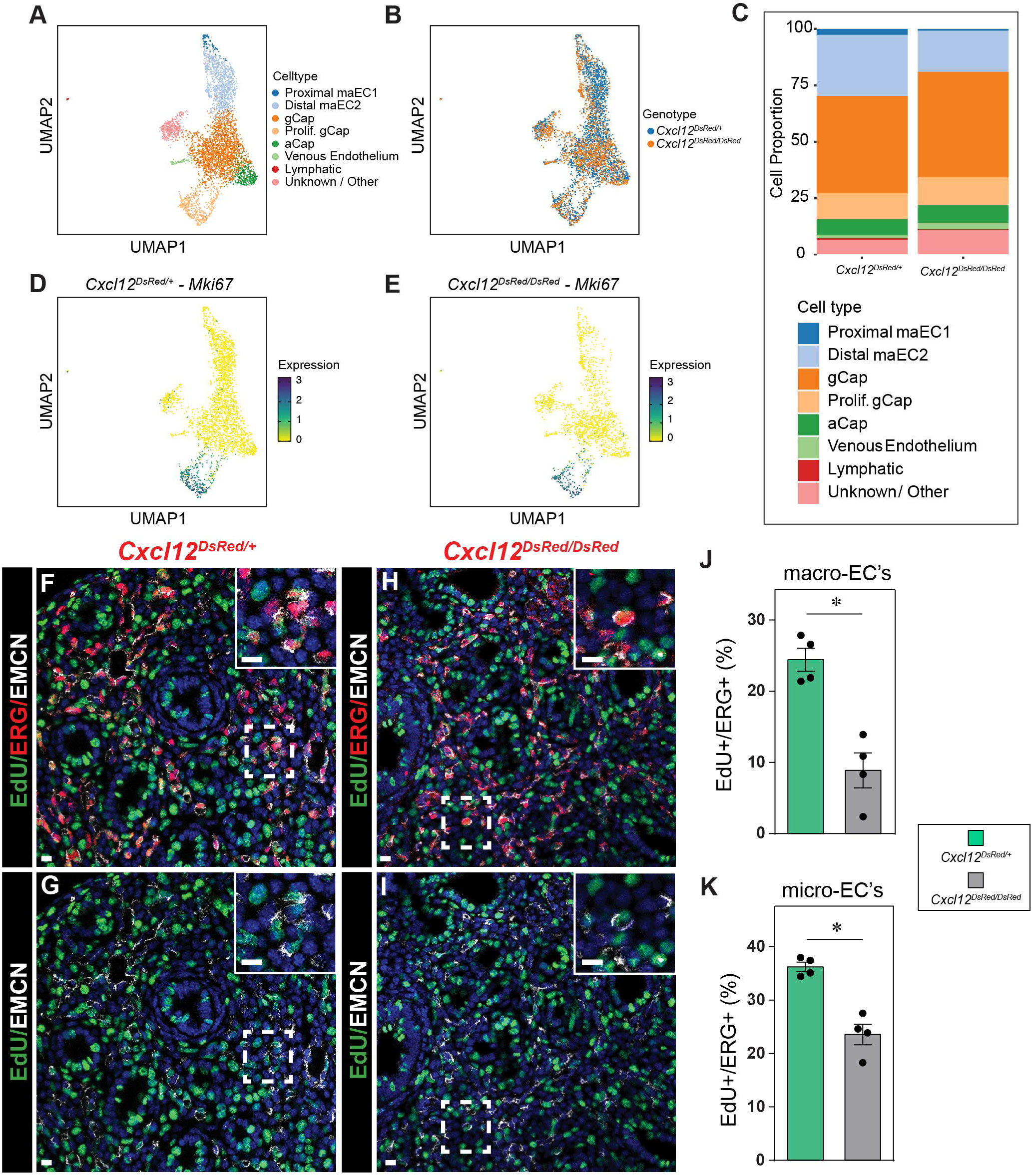
Decreased proliferation in control *Cxcl12*^*DsRed*^ versus null *Cxcl12*^*DsRed/DsRed*^ vasculature. (A) UMAP embedding of cells colored by endothelial cell populations at E18.5. (B) UMAP embedding of cells colored by *Cxcl12*^*DsRed/+*^ and *Cxcl12*^*DsRed/DsRed*^ ECs at E18.5. (C) Cell Proportion of *Cxcl12*^*DsRed/+*^ and *Cxcl12*^*DsRed/DsRed*^ at E18.5. (D,E) UMAP embedding of cells colored by *Mki67* expression in *Cxcl12*^*DsRed/+*^(D) and *Cxcl12*^*DsRed/DsRed*^ (E) ECs at E18.5. (F-K) Assement of proliferation measured by EdU incorporation in *Cxcl12*^*DsRed/+*^ and *Cxcl12*^*DsRed/DsRed*^ macro- and micro-ECs. (F-I) IHC for EdU, ERG, and EMCN to mark proliferating macro- and micro-ECs in in *Cxcl12*^*DsRed/+*^ (F,G) and *Cxcl12*^*DsRed/DsRed*^ (H,I) lung sections. (J,K) Quantification of percent EdU+/ERG+ macro-(J) and micro-ECs in *Cxcl12*^*DsRed/+*^ and *Cxcl12*^*DsRed/DsRed*^ lungs (n=4), *p<0.005. Scale Bar: F-I = 10 µm, insets = 10 µm.

To determine whether differences in proliferation are a mechanism responsible for the diminished number of ECs and pulmonary hypoplasia, we examined *Mki67* expression, noting no significant changes in the number of proliferating ECs at E18.5 by scRNA-seq (Fig 6C-E). We further interrogated proliferation by IHC at E15.5, an earlier time point when the vasculature demonstrates more rapid growth. Using antibodies to EMCN to mark capillary and venous endothelium, ERG for pan-endothelium, and 5-ethynyl-2’
s-deoxyuridine (EdU) incorporation for proliferation, we quantified the number of proliferating endothelial cells (Fig. 6F-K). At E15.5, we observed a significant decrease in the proliferation of both proximal macrovessel and distal plexus ECs in *Cxcl12*^*DsRed/DsRed*^ embryonic lungs, indicating an early proliferative defect in *Cxcl12*^*DsRed/DsRed*^ embryonic lungs.

### Loss of CXCL12 leads to significant transcriptome changes in the endothelial niche

Lung development is a coordinated process involving communication between cellular and tissue types. Failure of this reciprocal behavior results in transcriptional changes and defects in development, and this process can be elegantly annotated using scRNA-seq. Analysis of UMAP embedding of cells from two E18.5 control and two *Cxcl12*^*DsRed/DsRed*^ embryonic lungs revealed transcriptional changes in multiple endothelial niche compartments (Fig. S6A,B, Table S3). While overall numbers of cell populations were mostly unaffected, there were significant differences in cell type-specific transcriptomes in control versus null embryos. In particular, the immune cells, including the interstitial and alveolar macrophages, demonstrated significant changes in gene enrichment. In addition, the examination of the epithelium, including secretory, ciliated, alveolar type 1 (AT1), and alveolar type 2 (AT2) cells, revealed significant differences in gene expression (Figure S6E). Evaluating broad changes using the top upregulated and downregulated genes in knock out versus control whole lungs was performed with GO term enrichment. We observed that there was an enrichment of genes associated with branching involved in blood vessel morphogenesis and categories associated with mitochondrial bioenergetics in endothelial cells. The epithelium had enrichment of genes involved in the regulation of endothelin production. Immune cells expressed genes involved in microglial cell or macrophage activation, and mesenchymal cells were enriched in genes involved in membrane transport. (Figure S6D-G, Table S4).

## DISCUSSION

We have developed a temporal and spatially integrated transcriptomic atlas of *Cxcl12*+ endothelium during lung development. With these data, we identified two novel arterial endothelial cells populations in the developing lung (Fig. 7). This is one of the first studies uncovering functionally and spatially defined subpopulations in developing arterial endothelium. This distinction improves recognition of distinct roles for arterial ECs in patterning and assembly of the developing artery. In addition, we have uncovered a CXCL12 signaling gradient from the artery to the capillary network. The unique expression characteristics of CXCL12 receptors suggest haptotaxis as a possible mechanism for artery assembly in the lung. Finally, we have provided a framework for analysis of branching abnormalities in the pulmonary vasculature and uncovered several novel roles for CXCL12 in pulmonary vascular development, identifying a role in proliferation and directional migration for the lung.

**Figure 7.**
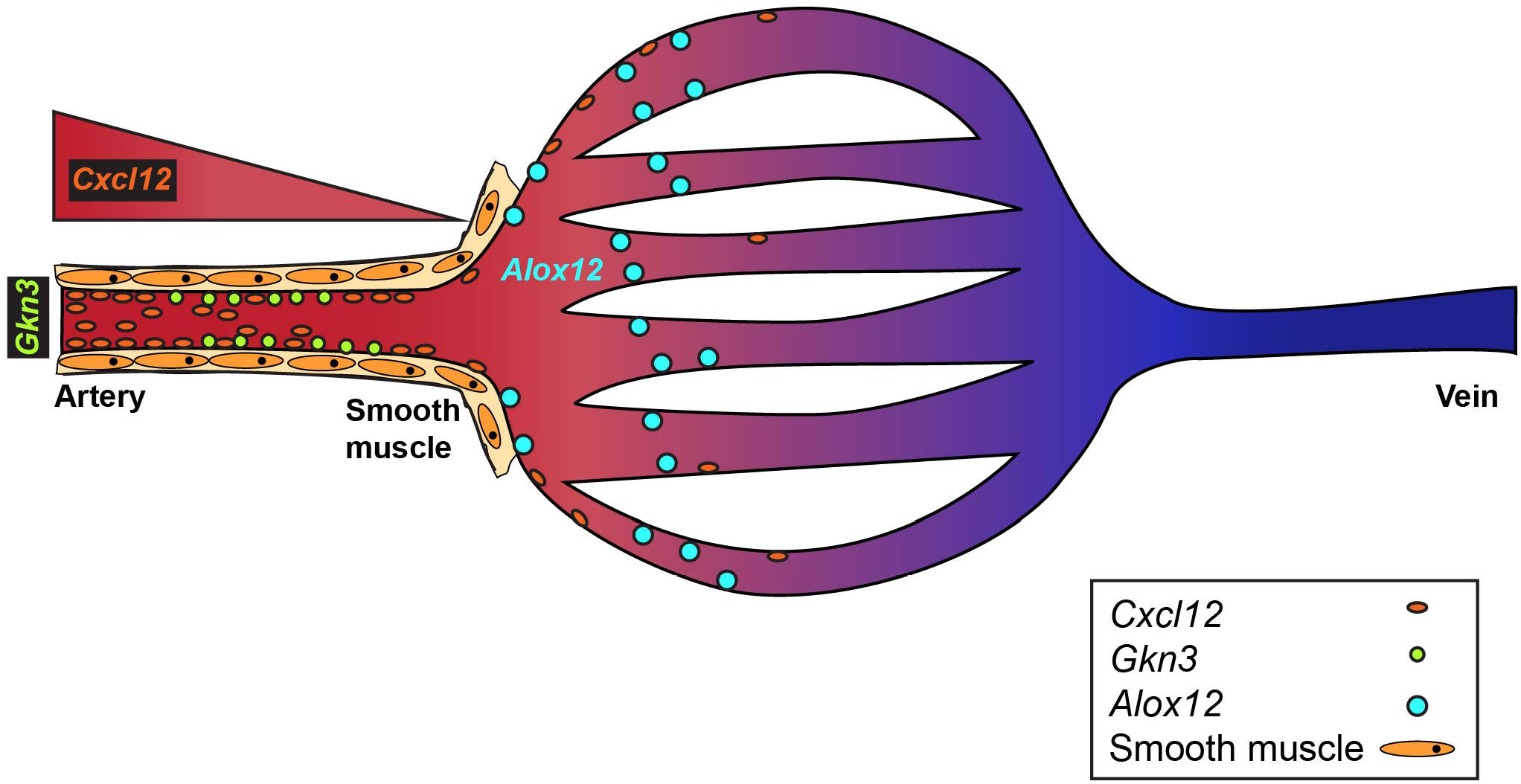
A model of CXCL12 signaling and the developing pulmonary vasculature.

### Proximal-distal patterning of the pulmonary artery indicates distinct spatial functions

From E15.5 to P8, the intrapulmonary artery segregates into proximal and distal subpopulations. Gene set enrichment on GO terms in these populations indicates distinct functions. The proximal artery expresses genes associated with extracellular structure and matrix formation, suggesting a role for vessel wall construction. By contrast, the distal artery comprises genes important in vascular development and morphogenesis. Previous descriptions of distinct subpopulations of arterial endothelium in other developing organs such as the coronary arterial system are limited to only subtypes of varying maturation stages (Su et al., 2018). While compartmentalization of the capillary endothelium of the lung has been previously demonstrated, this is one of the first studies defining the developing arterial endothelium into spatial and functionally different subpopulations in any organ system (Gillich et al., 2020; Niethamer et al., 2020; Vila Ellis et al., 2020).

Functionally distinct arterial endothelial subpopulations exist in other adult organs as well. The kidney contains 24 subpopulations of endothelium, and arterial endothelium can be divided into efferent, afferent, and juxtaglomerular apparatus-associated afferent arteriole populations by gene expression and function (Dumas et al., 2020). These 3 populations have previously established physiological functions within the kidney. Similarly, the aorta is segregated into subpopulations that regulate different arterial responses at homeostasis and hypertensive states in humans and mice (Kalluri et al., 2019; Zhang et al., 2021). Similar findings exist in a study on the adult human pulmonary artery endothelium in health and disease. The pulmonary artery endothelium has been profiled from adult control and patients with pulmonary arterial hypertension (PAH) (Asosingh et al., 2021). In this study, scRNA-seq based clustering uncovered heterogeneity including arterial endothelial cell subpopulations enriched for genes of quiescence, proliferation, and angiogenesis. PAH subpopulations demonstrated a higher contribution of cells with gene enrichment for angiogenesis and proliferation, consistent with known pathological findings in PAH. Whether clusters enriched for angiogenesis corroborate our developmental arterial clusters is unknown. For this investigation, cells were only isolated from the main PA and 1^st^ through 4^th^ branches whereas our data represent more distal and terminal branches of the developing arterial tree. Nevertheless, comparison of top genes in the human control and PAH angiogenic clusters with our data do suggest *CXCR4* as a shared gene. *Cxcr4* is enriched in our distal angiogenesis cluster and suggests a common pathway in development and disease.

Our unbiased cell clustering and gene annotation of these clusters identified *Gkn3* and A*lox12* as most representative of the proximal and distal maECS, respectively (Fig. 7). GKN3 was previously identified as an arterial endothelial marker in the brain (Vanlandewijck et al., 2018), indicating its potential as a marker for subpopulations of the arterial endothelium in other vascular beds. ALOX12 has been implicated in the regulation of angiogenesis in other organs and maladaptive vascular remodeling in pulmonary hypertension (Preston et al., 2006; Zhang et al., 2018; Zheng et al., 2020). This functional prediction is in line with considering ALOX12 as a marker for distal artery endothelium. Similarly, GO analysis of proximal artery endothelium suggested a role in extracellular structure and matrix formation. The importance of our ability to distinguish functional endothelial cellular compartments is two-fold. First, we have highlighted genes critical for angiogenesis that are distinct from those involved in vascular wall assembly. This distinction provides the framework for studying known and novel signaling pathways in a compartmental and functional process, to identify potential mechanisms and therapies for vascular regeneration. Second, components of signaling pathways such as Notch signaling are broadly expressed in arterial EC populations (Mack and Iruela-Arispe, 2018). Our arterial EC subpopulation gene enrichment analysis now allows us to define differential expression of receptors, ligands, and/or transcription factors in these specific arterial subpopulations to improve interrogation of signaling mechanisms. As such, future studies could be designed to examine the interactions of both distal and proximal arterial endothelia within their specific niches during vascular growth and assembly.

### CXCL12 signaling defines lung EC compartments and regulates vascular growth

CXCL12 signaling is an evolutionarily conserved angiogenic pathway important for vascular development and regeneration (Red-Horse and Siekmann, 2019). The pathway functions primarily in the recruitment of cells via chemotaxis-directed migration, but it is also implicated in cellular proliferation (Bianchi and Mezzapelle, 2020; Majumdar et al., 2014). Directed cell migration is mediated by a gradient of CXCL12 expression with cellular recruitment to the area of highest expression. One example is the coronary arterial system derived from a vascular plexus that arises from the sinus venosus (Red-Horse et al., 2010). Disruption of this gradient by deletion of CXCL12 results in failure of the coronary artery system to connect to the aorta (Ivins et al., 2015). Although the connection of the pulmonary artery to the heart is intact in our knockout, aberrant branching and growth of the pulmonary arterial tree has been described after disruption of the CXCL12 gradient (Kim et al., 2017). We have further expanded on this finding and have quantified branching defects seen in *Cxcl12*^*DsRed/DsRed*^ mutants. Furthermore, we discovered a role for CXCL12 in growth of the pulmonary vasculature. Our mutants have significant reduction in arterial, venous, and capillary density, suggesting ubiquitous pulmonary vascular hypoplasia. Capillary EC deficits have been previously described in an adult pneumonectomy regeneration model in mice. Deletion of CXCL12 in platelets or CXCR4 alone or combined with ACKR3 deletion resulted in a significant reduction in pulmonary capillary EC proliferation, indicating fetal programs as a mechanism for regeneration in the lung (Rafii et al., 2015). Indeed, a CXCL12 therapeutic approach has been used to treat hyperoxia-induced neonatal rodent lung injury (De Paepe et al., 2020; Guerra et al., 2019).

The CXCL12 signaling axis is expressed in a compartmental fashion. There is a morphogen gradient of CXCL12 highest in the arterial ECs with diminution of expression in the capillary ECs. CXCL12 is also expressed at lower levels in the vascular smooth muscle and adventitial fibroblasts. While this study did not assess endothelial or mesenchymal-specific deletion of CXCl12, the majority of CXCL12 is expressed in ECs (Negretti et al., 2021; Zepp et al., 2021). In addition, expression of the receptors for CXCL12 indicates an endothelial-dependent signaling mechanism for vascular assembly and growth. Indeed, deletion of CXCR4 using a pan-endothelial Cre recombinase corroborated the branching defects seen with global CXCL12 deletion (Kim et al., 2017). CXCR4 is found on the distal artery and capillary ECs, and ACKR3 is expressed in capillary and venous endothelium. This suggests directional migration of CXCR4 positive endothelium into the growing arterial tree as a mechanism of arterial assembly and growth, a process first described in zebrafish fin regeneration (Xu et al., 2014). In addition, CXCR4 and ACKR3 expression in their respective compartments could explain the reduction in capillary and venous density observed in *Cxcl12*^*DsRed/DsRed*^ mutants. Future studies deleting these receptors from arterial, capillary, or venous endothelium should clarify these mechanisms.

### The CXCL12 endothelial signaling niche affects multiple cell types in the lung

UMAP-based clustering and cell annotation in our scRNA-seq on control versus *Cxcl12*^*DsRed/DsRed*^ mutant embryonic lungs revealed significant cellular population and transcriptomic changes in multiple cell types. ECs and mesenchyme-associated niche cells (MANCs) appeared to be reduced in number. The EC affect was pan-vascular and represents our similar findings on density quantifications. The transcriptomic differences occurred in several tissue compartments. In particular, the immune system and epithelium lacked integration in our clustering, suggesting important changes in the immune cells and epithelium.

For the immune system, CXCL12 and its chemotactic ability for immune cells is well studied. Expression of CXCL12 in the endothelium allows recruitment and transmigration of cells across the vessel wall to areas of infection and inflammation (Cheng et al., 2014). In our data, transcriptomic changes appeared predominantly in alveolar and interstitial macrophages. CXCR4 is broadly expressed by immune cells in our dataset, but ACKR3 is restricted to macrophages. Both receptors are implicated in recruitment of macrophages and differentiation from monocytes (Man et al., 2012; Sanchez-Martin et al., 2011). Despite this evidence, it is not known whether this is an early developmental defect arising in the bone marrow or innate to the lung given we are using a global knock out of CXCL12.

Epithelial-endothelial interactions play an important role in lung development and regeneration. Growth factors such as vascular endothelial growth factor A (VEGFA), Wingless and Int-1 (WNT) ligands, and sonic hedgehog (SHH) mediate reciprocal communication to orchestrate both airway and vascular development (Cornett et al., 2013; Del Moral et al., 2006; Peng et al., 2013; Vila Ellis et al., 2020; Yamamoto et al., 2007). While these interactions are direct in lung development, the mechanism of CXCL12 signaling mediating vascular development is unclear. In our single-cell profiling of lung cells, the epithelium does not express CXCR4, ACKR3 or any additional chemokine receptors in this family. This may be a result of the limitations of scRNA-seq and its sequencing depth as there is other evidence of CXCR4 expression in AT2s in humans (Murdoch et al., 1999). Alternatively, there may be an indirect relationship mediated via an additional niche cell such as mesenchyme or immune cell populations.

In summary, we have defined a CXCL12 signaling single-cell atlas. In doing so, we uncovered arterial heterogeneity with distinct functional transcriptomes. These findings will improve the clarity of signaling mechanisms along the pulmonary arterial tree. In addition, the CXC12 signaling axis plays an important role in all developing lung endothelial compartments. As such, CXCL12 signaling may prove a therapeutic target for diverse forms of pulmonary vascular hypoplasia.

## MATERIALS AND METHODS

### Animals

Development and genotyping information of mouse line *Cxcl12*^tm2.1Sjm/^J (Jackson Laboratories, Stock no.: 022458, Cxcl12^DsRed^) has been previously described (Ding and Morrison, 2013). Timed matings were performed counting day of plug as day 0.5. Lungs were harvested from embyros from both heterozygous (*Cxcl12*^*DsRed/+*^) and homozygous knockout (*Cxcl12*^*Dsred/DsRed*^) mice, and they were prepared as noted below for whole mount and tissue section staining. All animal studies were performed in strict adherence to the guidelines of the Children’s Hospital of Philadelphia Institutional Animal Care and Use Committee.

### Whole mount imaging

Lungs were harvested at E12.5, E13.5, E15.5, E17.5 and E18.5 from both heterozygous (*Cxcl12*^*DsRed/+*^) and homozygous knockout (*Cxcl12*^*Dsred/DsRed*^) mice. Based on previously established protocols (Das et al., 2019), lungs were fixed in 2% paraformaldehyde for 4 hours and then imaged for DsRed to evaluate the arterial endothelium or prepared for immunofluorescence-based whole mount imaging. Lung lobes were incubated with primary antibody (anti-VWF, rabbit, 1:100, F3520, Sigma and anti-EMCN, goat, 1:200, AF4666, R&D Systems), in 5% donkey serum and 0.5% Triton X-100 in phosphate buffered saline (PBST) for 4 hours at room temperature (RT) on a shaker, and overnight (O/N) at 4°C on a shaker. The samples were washed in 0.5% PBST (4 × 30 minutes at RT), O/N at 4°C on a shaker. The samples were then incubated in secondary antibody (donkey anti-mouse Alexa Fluor 488, A-21202 and anti-goat Alexa Fluor 647, A21447, ThermoFisher, 1:250), in 5% donkey serum and 0.5% PBST for 1 hour at RT on a shaker followed by O/N at 4°C. The lungs were washed in 0.5% PBST (4 × 30 minutes at RT), fixed in 2% paraformaldehyde for 2 hours at RT and then washed with PBS (2 × 30 minutes at RT). Finally, the lungs were cleared using CUBIC (RIA reagent) O/N at 37°C, and then stored at 4°C until imaged using a Leica DMi8 Thunder system and/or Leica DMi8 confocal microscope.

### Immunohistochemistry

Lungs were harvested and fixed as previously described(Frank et al., 2019). They were then washed, dehydrated, embedded in paraffin, and 6-μm sections were obtained. IHC was performed on E15.5, E17.5 and E18.5 sections. In brief, the sections were deparaffinized and rehydrated followed by antigen retrieval (Reveal decloaker, MSPP-RV1000M, Biocare Medical). The sections were incubated with 3% H_2_O_2_ for 15 minutes to quench endogenous peroxidases, blocked with 5% donkey serum, and finally, incubated with primary antibody in 0.1% PBST (VWF, EMCN, and RFP, rabbit, 1:50, 600-401-379, Rockland Immunochemicals) O/N at 4°C. The presence of relevant proteins was visualized using Alexa Fluor secondary antibodies (donkey anti-mouse AF488, anti-rabbit AF555, A-31572, Thermofisher and anti-goat AF647).

### RNA in situ hybridization

RNA ISH was performed with the RNAscope™ Multiplex fluorescent V2 Assay (323100, Advanced Cell Diagnostics (ACDbio), Newark, CA) per the manufacture’s recommendations. RNAscope 3-plex negative control probe (*DapB*) and mouse specific 3-plex positive control probe (*Polr2a*) were used for control sections. RNAScope target probes include *Cxcl12, Cxcr4, Ackr3, Gkn3, Alox12, Gja5, Nrg1, Apln*, and *Gpihbp1*, and images were obtained on channels C1, C2, C3 and C4 with Opal fluorophore reagents (Opal 520 FP1487001, Opal 540 FP1494001, Opal 570 FP1488001 or Opal 650 FP1496001, Akoya Biosciences). Finally, slides were incubated in DAPI for 30 seconds and then mounted using Prolong Gold Antifade Mountant (P36930, ThermoFisher).

### Quantification of vascular density and vein counts

RNA ISH combined with IHC was performed on sections as noted above. Capillary and arterial density were measured using Imaris software (version 9.7.2) on 40X magnification confocal z-stacks. From whole lung, 6 identical regions were selected per mice lung. For each image, the region of whole tissue was measured with 3D reconstruction of DAPI stained tissue using surface function. The total tissue volume was acquired via volume statistics function in Imaris. In order to measure vessel density, a single channel of macrovascular (VWF), capillary (EMCN) or arterial marker (Cxcl12-DsRed) was also reconstructed in 3D using surface function. For capillary density, macrovasculature (VWF) was excluded from 3D reconstruction to measure only capillaries. The tissue volume was measured using volume statistics. Same threshold parameters were applied when comparing both knock out and control groups. Both capillary and arterial density were obtained by dividing the capillary or arterial volume by the total lung volume per image (Chen et al., 2021; Nizari et al., 2019).

Vein count was measured from 40X magnification confocal images, with 6 identical regions selected per mice lung. For each image, vessels stained with an arterial marker (*Gja5* RNA scope probe or DsRed) and a macrovascular endothelial marker (VWF) were manually counted. The final vein count was obtained by excluding arteries from the total number of macrovessels stained. The vein count was then divided by the total number of vessels to avoid bias of image selection.

### Quantification of proliferation

To quantify the proliferative endothelial cells, we first performed IHC for pan-endothelium (anti-ERG, rabbit, 1:100, ab214341, Abcam) and EMCN that stains for capillary and venous endothelium. Following IHC, we performed a click chemistry-based EdU proliferation assay (Click-iT™ Plus EdU Alexa Fluor™ 647, C10640, ThermoFisher). We captured 5 random areas on each tissue section from each embryo. We counted the percentage of KI67+ cells in arteries, veins and capillaries. Briefly, we counted all ERG+ cells to determine the total endothelial cells in the field of view. We classified EMCN-/ERG+ cells as arterial endothelial cells and EMCN+/ERG+ cells as either capillary or venous endothelial cells. We differentiated venous cells from capillary cells by tracing a subset of cells which formed a lumen. Among these, KI67+ cells were counted to quantify percentage of proliferative cells in each subtype.

### Endothelial cell isolation

For the scRNA-seq time course of CXCL12+ endothelium, *Cxcl12*^*DsRed/+*^ embryos were harvested at E13.5, E15.5, E18.5, and P8. For control versus knock out lung scRNA-seq comparison, *Cxcl12*^*DsRed/+*^ and *Cxcl12*^*DsRed/DsRed*^ embryos at E18.5 were harvested. Lungs were removed, minced and processed into a single cell suspension in a Collagenase I (8 mg/5ml, 17100017, Life Technologies) /Dispase (1:10, 354235, Corning)/RNase-free DNase (1:500, M6101,Promega) solution for 20 min at 37°C (Frank et al., 2016). CD31+TdTomato+ cells underwent FACS-based isolation from single cell suspension using a MoFlo Astrios EQ (Beckman Coulter) flow cytometer with antibody staining for CD31-PECy7 (25-311-82, ThermoFisher), CD45-APC (17-0451-82, ThermoFisher), and EpCAM-FITC (11-5791-82, ThermoFisher). Following negative selection for CD45 and EpCAM, CD31+/TdTomato+ ECs and CD31-/TdTomato+ mesenchymal-enriched cells were collected in FACS buffer without EDTA.

### scRNA-seq library preparation and next-generation sequencing

Cells were pelleted and live cell concentration was determined using trypan blue staining. The cell pellet was resuspended as per the 10X Genomics recommendation, and 16,500 cells were loaded onto the 10X Chromium (10X Genomics), to target a recovery of 10000 cells. The libraries were prepared according to the manufacture’s protocol using 10x Single cell 3’ v3 chemistry. These libraries were then sequenced on Illumina HiSeq 2500 instrument. CellRanger Count v3.1 (10X Genomics) was used to align reads onto the mm10 reference genome.

### Analysis of single cell sequencing data

For the *Cxcl12* knockout dataset, but not the dataset of *Cxcl12+* cells only, ambient background RNA were cleaned from the scRNA-seq data with “SoupX” as described previously (Schuler et al., 2021; Young and Behjati, 2020). The following genes were used to estimate the non-expressing cells, calculate the contamination fraction, and adjust the gene expression counts: *Dcn, Bgn, Aspn, Ecm2, Fos, Hbb-bs, Hbb-bt, Hba-a1, Hba-a2, Lyz1, Lyz2, Mgp, Postn, Scgb1a1*. For all datasets, quality filtering was then used to remove cells with > 10% or < 0.5% mitochondrial mRNA, and to remove cells with < 700 detected genes.

Dimensionality reduction, clustering, and visualization was performed using Seurat v3.2.2 and SCTransform (Hafemeister and Satija, 2019; Stuart et al., 2019), with manual inspection of the expression patterns of the marker genes: *Pecam1, Ccl21a, Vwf, Nrg1, Plvap, Car4*, and *Mki67*. SCTransform was run with each sequencing reaction as a batch variable with the percentage of mitochondrial RNA as a regression variable. Further data cleaning was done to remove gene counts for *Gm42418*, which is likely a rRNA (Kimmel et al., 2020). After removal of *Gm42418*, genes with expression in less than 10 cells across the dataset were removed from further analysis.

SCTransform (version 0.3.2.9008) was used with glmGamPoi (version 1.2.0) to normalize the data prior to PCA, UMAP embedding, and cell clustering (Ahlmann-Eltze and Huber, 2021). For the Cxcl12+ sorted dataset, Louvain clustering was performed on the first 25 principal components with a resolution of 0.2. For the dataset investigating the effect of Cxcl12 knockout, Louvain clustering was performed on the first 20 principal components with a resolution of 0.8. The difference in resolution was due to the number of cell types present in these two different experiments. All charts and heatmaps as part of the scRNA-seq analysis were generated with ggplot2, and all parts of the analysis were run in R 4.0.2. PANTHER GO enrichment analysis (version 10.5281/zenodo.5725227 released 2020-11-01) was performed on the top 40 genes enriched for interrogated clusters and reported by -log10(FDR) (Ashburner et al., 2000; Gene Ontology, 2021; Mi et al., 2019).

Predictions of RNA velocity were made using a velocyto, scVelo, CellRank pipeline. Spliced and unspliced mRNA counts were determined with velocyto (version 0.17.17) (La Manno et al., 2018) and analyzed using scVelo (version 0.2.3) with the dynamical model (Bergen et al., 2020). CellRank (version 1.5.0) was then used to infer the initial and terminal cell states (Lange et al., 2022). All packages were run in Python 3.7.11. A complete collection of all package versions, and code for all steps of the analysis is available at https://github.com/SucreLab/Cxcl12LungVascularDevelopment

## Acknowledgements

We would like to thank the Flow Cytometry core and Florin Tuloc at CHOP for cell sorting and collection. We would like to acknowledge Jacqueline Smiler, Fernanda Mafra Thompson, Renata Pellegrino da Silva, Michael Gonzalez, and James Garifallou in the CHOP Center for Applied Genomics for assistance with scRNA-seq library preparation and sequencing. And we are grateful to the Ayla Gunner Prushansky Research Fund.

## Footnotes

### Data availability

All sequencing data has been deposited in GEO under pending accession numbers

## FIGURE LEGENDS

**Figure S1. *Cxcl12*-DsRed+ endothelium is arterial in nature. (A**,**B)** Whole mount imaging of lung using VWF, DsRed, and EMCN at E12.5 (A) and E13.5 (B). **(C-F)** IHC staining for ERG/DsRed/EMCN at E15.5 and E18.5. Scale bar: A-B = 500 µm, C-F = 50 µm, insets = 10 µm.

**Figure S2. Gene expression and correlation with *Cxcl12***. Heat map of highly correlated genes in all EC populations.

**Figure S3. Characterization of *Cxcl12*+ ECs in proximal and distal vasculature**. (A,B) IHC staining for ELN (tropoelastin) and SMA with RNA ISH for *Gkn*3 at E15.5 (A) and E18.5 (B). (C,D) RNA ISH for *Cxcl12, Apln*, and *Gpihbp1* gene expression in the capillaries at E18.5. * represents *Cxcl12* expression in capillaries. Scale bar: A-D = 10 µm, insets = 10 µm.

**Figure S4. Confirmation of *Cxcl12, Cxcr4*, and *Ackr3* expression in a whole lung scRNA-seq dataset**. (A) UMAP embedding of endothelial cells only colored by EC populations from publicly available published dataset. (B) UMAP embedding of cells colored by *Cxcr4* expression. (C) UMAP embedding of cells colored by *Ackr3* expression. (D) UMAP embedding of cells colored by *Cxcl12* expression. (E-G) Dot plots for *Cxcr4* (E), *Ackr3* (F), and *Cxcl12* (G) expression in EC populations at various time points. (H-K) Spatial transcriptomic assessment for Cxcr4 and Ackr3 expression in capillary and vein ECs. (H,I) RNA ISH for *Ackr3* and *Nrg1* (venous ECs) or Gpihbp1 (capillary ECs) at E18.5. (J,K) RNA ISH for *Cxcr4* and Aplnr (capillary ECs) at E18.5 or *Gja5* (arterial ECs). * represents capillary expression. Scale bar: H-K = 10 µm, insets = 10 µm.

**Figure S5. Loss of *Cxcl12* leads to aberrant branching and vascular hypoplasia**. (A-C) Assessment of peripheral artery branching of *Cxcl12*^*DsRed/+*^ (A)and *Cxcl12*^*DsRed/DsRed*^ (B) embryonic lungs. Insets: Higher magnification of distal arteries. (C) Comparative branching angle measurements of *Cxcl12*^*DsRed/+*^ versus *Cxcl12*^*DsRed/DsRed*^ arteries (n=4, Statistical significance: *p<0.05). (D) Sholls analysis showing distribution of each sample and (E) Scatter plot of the average Sholls intersections at all distance from the origin of artery in *Cxcl12*^*DsRed/+*^ versus *Cxcl12*^*DsRed/DsRed*^ embryonic lungs (n=4, Statistical significance: *p<0.05). (F-H) Representative volumetric analysis of capillaries, veins and arteries using Imaris 3D surface rendering. (F) Volumetric analysis of capillaries in *Cxcl12*^*DsRed/+*^ (upper panel) versus *Cxcl12*^*DsRed/DsRed*^ embryonic lungs (bottom panel). (G) Volumetric analysis of veins in *Cxcl12*^*DsRed/+*^ (upper panel) versus *Cxcl12*^*DsRed/DsRed*^ embryonic lungs (bottom panel). (H) Volumetric analysis of arteries in *Cxcl12*^*DsRed/+*^ (upper panel) versus *Cxcl12*^*DsRed/DsRed*^ embryonic lungs (bottom panel). Arterial and Venous percentage in *Cxcl12*^*DsRed/+*^ versus *Cxcl12*^*DsRed/DsRed*^ embryonic lungs (n≥4, Statistical significance: *p<0.05). Scale bar: A-B = 500 µm, Insets = 50 µm F-H = 50 µm.

**Figure S6. scRNA-seq analysis reveals global transcriptome changes in *Cxcl12***^***DsRed/DsRed***^ **whole lungs**. (A) Integrated *Cxcl12*^*DsRed/+*^ and *Cxcl12*^*DsRed/DsRed*^ UMAP embedding of cells colored by cell type to compare *Cxcl12*^*DsRed/+*^ versus *Cxcl12*^*DsRed/DsRed*^ embryos at E18.5. (B) UMAP embedding of whole lung cells colored by genotype to compare *Cxcl12*^*DsRed/+*^ versus *Cxcl12*^*DsRed/DsRed*^ embryos at E18.5. (C) Changes in cell type proportion between *Cxcl12*^*DsRed/+*^ and *Cxcl12*^*DsRed/DsRed*^ embryonic lungs. (D-G) PANTHER GO analysis for top 5 ranked categories using either upregulated (UP) or downregulated gene enrichment sets in cell populations. (D) GO analysis for ECs. (E) GO analysis of epithelial cells. (F) GO analysis of immune cells. (G) GO analysis of mesenchymal cells.

