## Supplementary figures and images for "A CXCL12 morphogen gradient uncovers lung endothelial heterogeneity and promotes distal vascular growth"

### Supplmental Figure 1

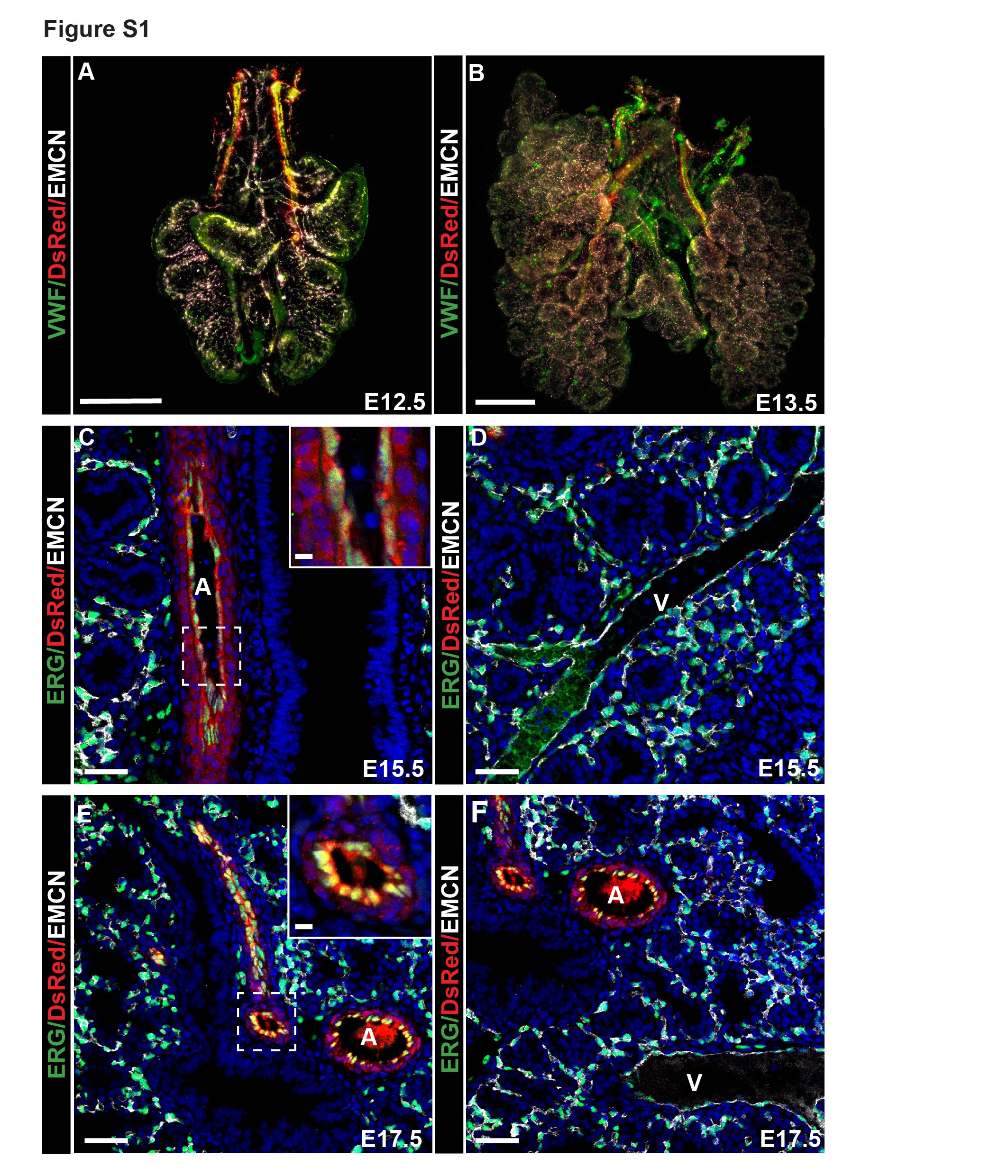

### Supplmental Figure 2

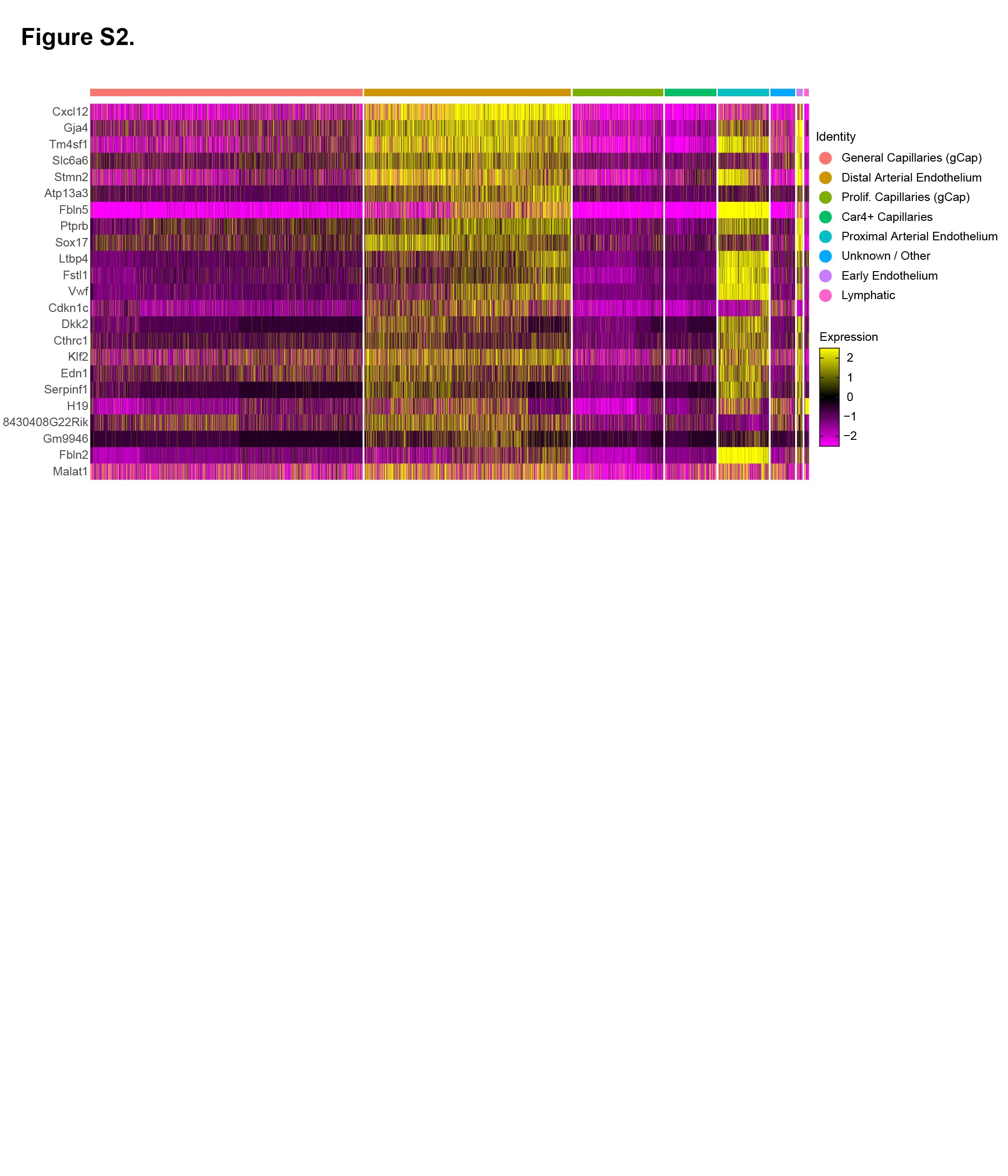

### Supplmental Figure 3

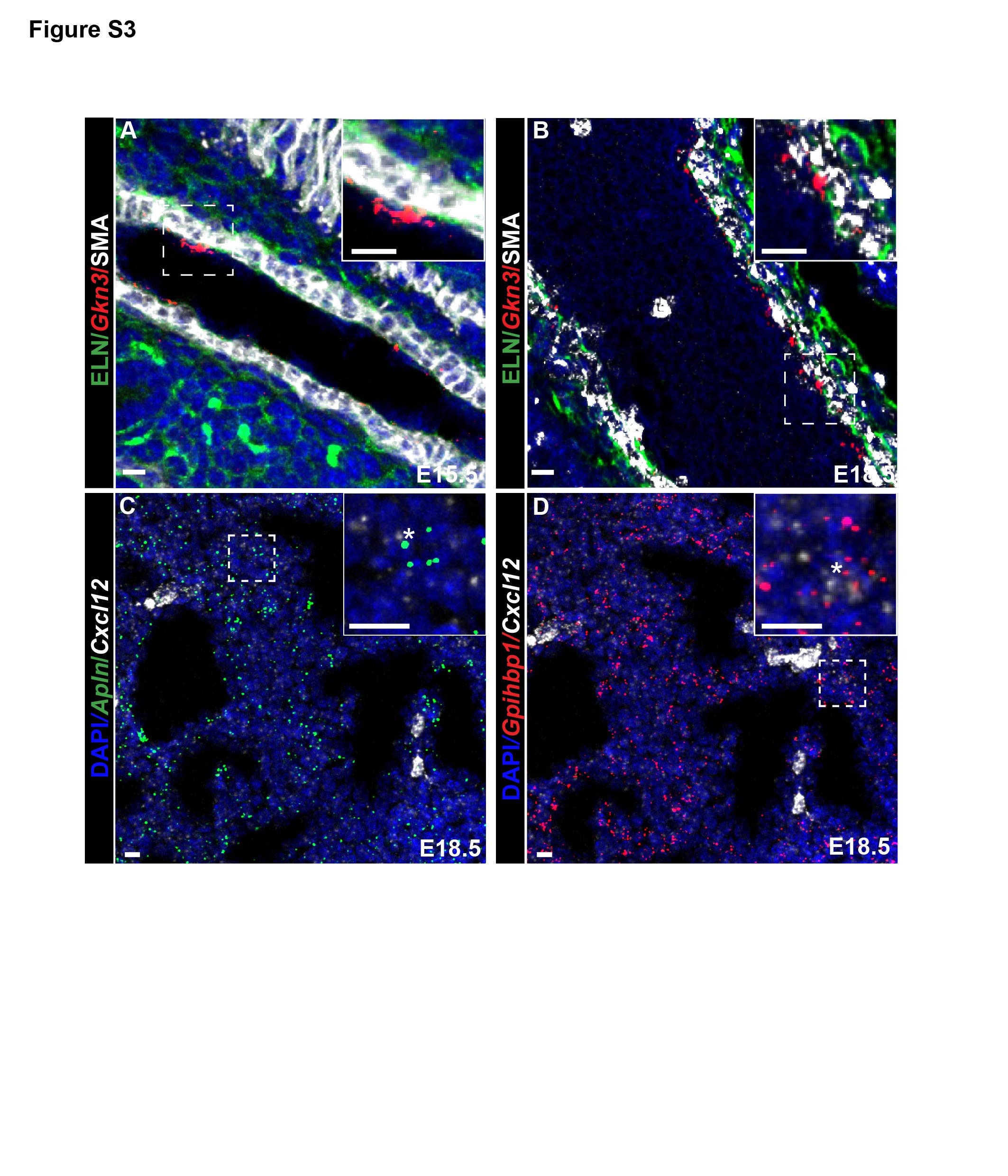

### Supplmental Figure 4

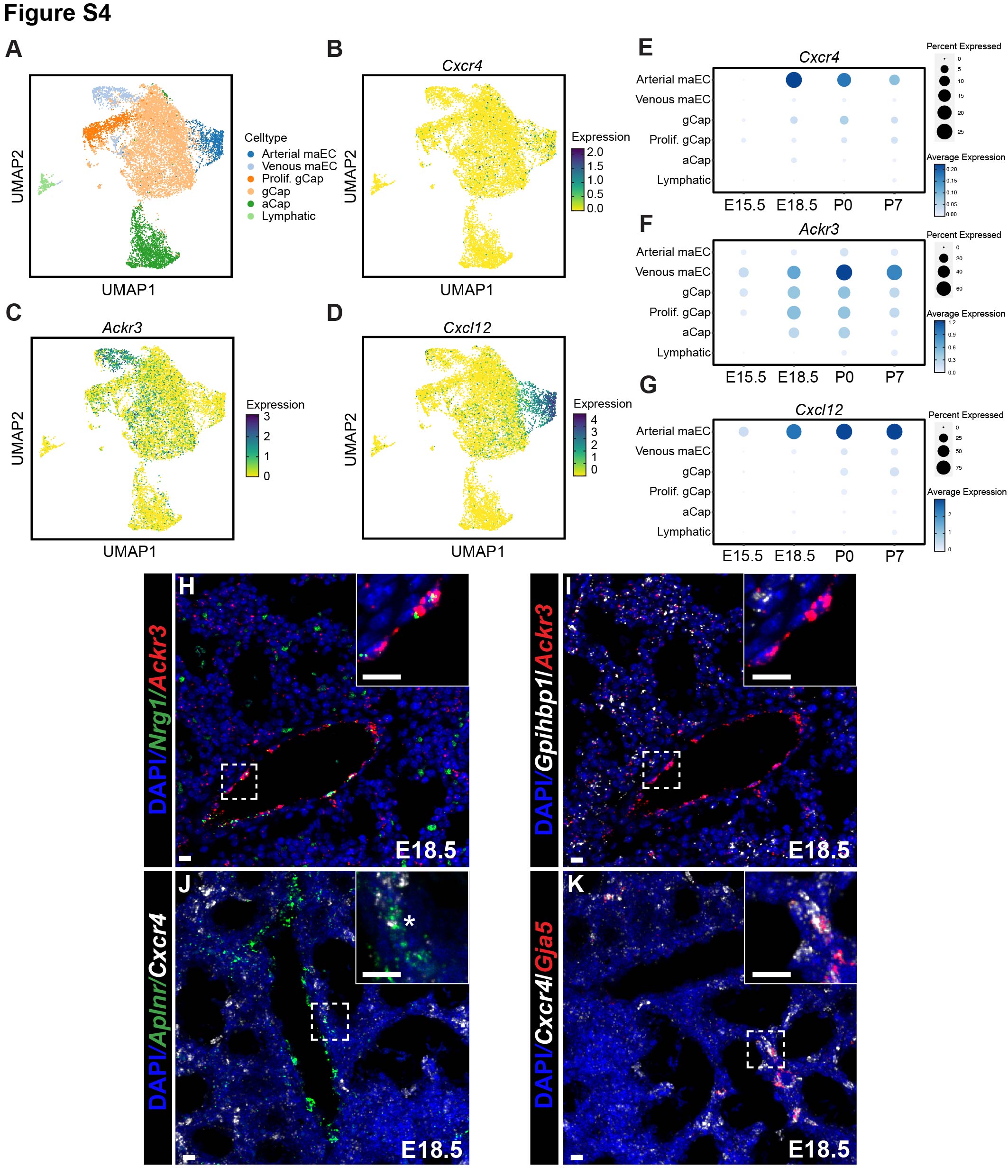

### Supplmental Figure 5

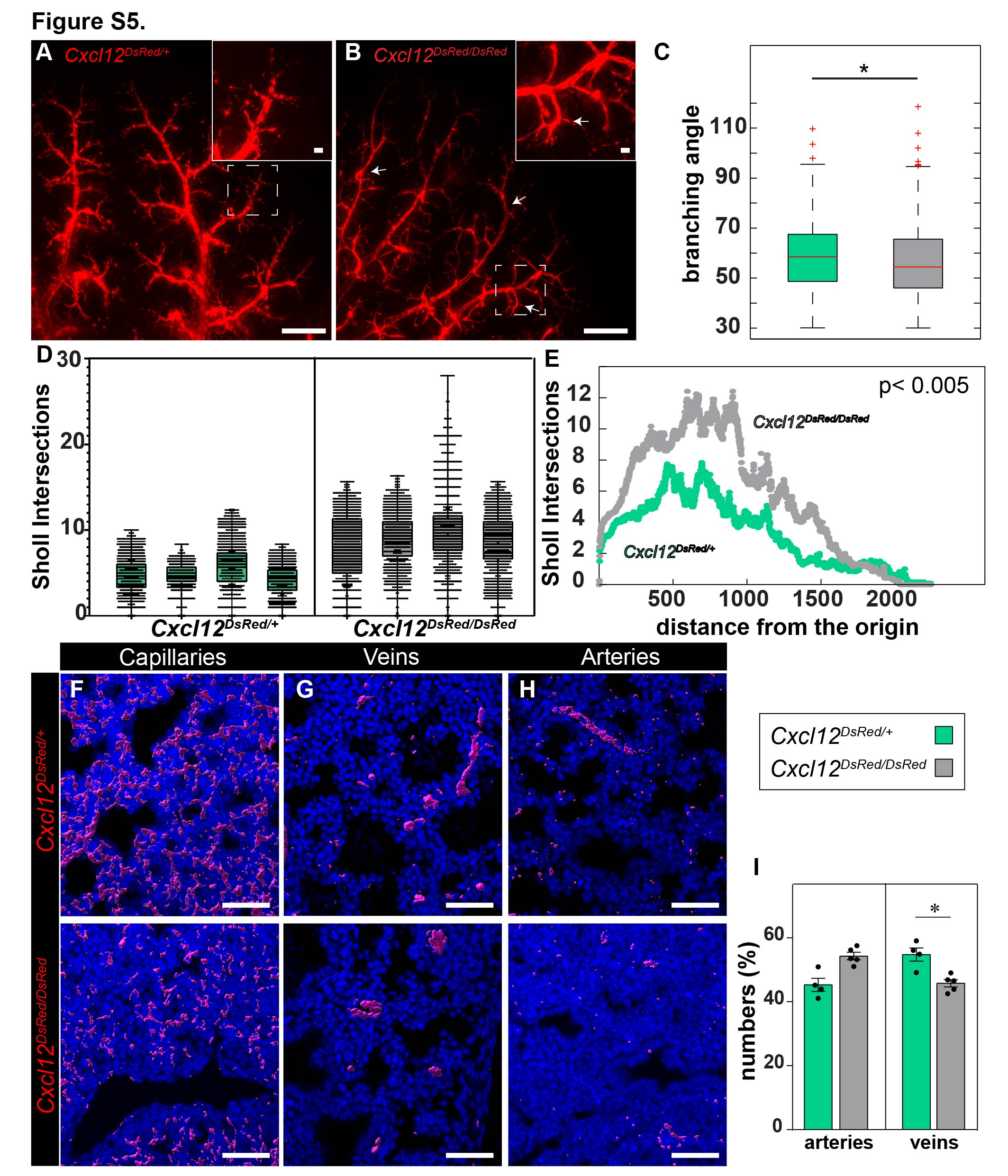

### Supplmental Figure 6

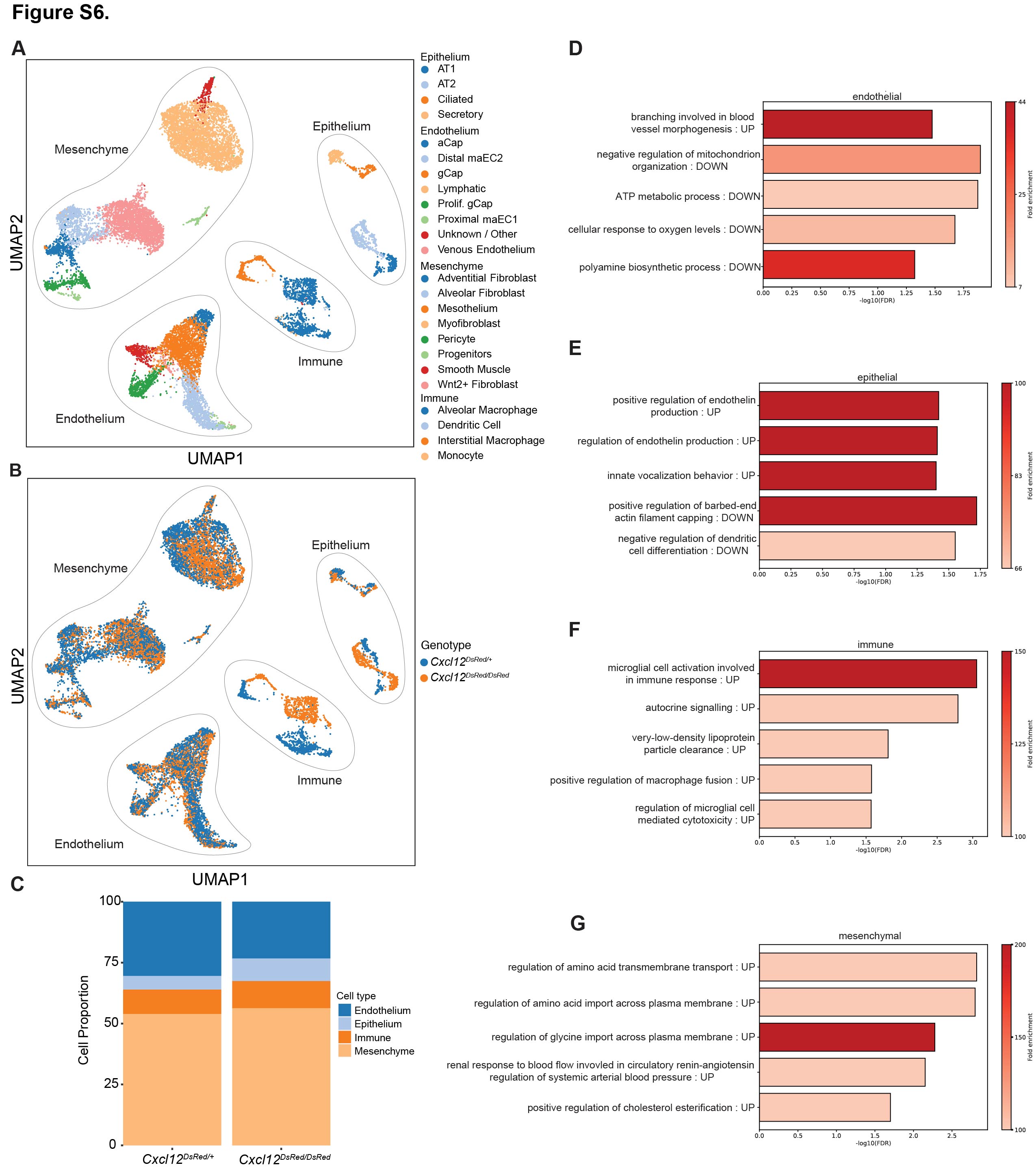
